# Risk factors associated with Avian Influenza subtype H9 outbreaks on poultry farms in Kathmandu valley, Nepal

**DOI:** 10.1101/782375

**Authors:** Tulsi Ram Gompo, Bikash Raj Shah, Surendra Karki, Pragya Koirala, Manju Maharjan, Diker Dev Bhatt

## Abstract

Poultry sector contributes to four percent in national GDP of Nepal. However, this sector is under threat with periodic outbreaks of Avian Influenza (AI) subtypes H5 and H9 since 2009. This has been both a both public health threat and an economic issue. Since last three years, outbreaks of AI subtype H9 has caused huge economic losses in major poultry producing areas of Nepal. However, the risk factors associated with these outbreaks have not been assessed. A retrospective case-control study was conducted from April 2018 to May 2019 in Kathmandu Valley to understand the risk factors associated with AI subtype H9 outbreaks. Out of 100 farms selected, 50 were “case” farms, confirmed positive to H9 at Central Veterinary Laboratory, Kathmandu, and other 50 farms were “control” farms, matched for farm size and locality within a radius of three km from the case farm. Each farm was visited to collect information using semi-structured questionnaire. Nineteen potential risk factors were included in the questionnaire under the broad categories: birds and farm characteristics, management aspects and biosecurity status of the farms. Univariable and multivariable logistic regression analysis were conducted to calculate corresponding odds ratios. Identified risk factors associated with AI subtype H9 outbreaks in Kathmandu valley were: “Birds of age 31-40 days” (OR= 11.31, 95% CI: 1.31-98.02, p=0.028), “Older farms operating for >5 years” (OR= 10.9, 95% CI: 1.76-66.93, p=0.01), “Commercial layers farms” (OR=36.0, 95% CI: 0.97-1332.40, p=0.052), “Used stream water to water birds (OR= 5.7, 95% CI: 1.10-30.13, p=0.039)”, “Farms without practice of fumigation after each batch of poultry (OR= 4, 95% CI: 1.44-13.13, p=0.009)., “Farm with previous history of AI (OR= 13.8, 95% CI: 1.34-143.63, p = 0.028), “Did not applied farm boots (OR= 2.58, 95% CI: 0.98-6.80, p= 0.055), “Visitors allowed to enter the farms (OR= 2.5, 95% CI: 1.011-6.17, p = 0.047) and “No foot bath at entry of farms (OR= 3.3, 95% CI: 1.29-8.38, p = 0.013). This study depicts that outbreaks of AI subtype H9 in Kathmandu valley was related to poor management practices and biosecurity in the poultry farms. We suggest improving management practices and increase biosecurity in the farms to reduce incidences of AI subtype H9 outbreaks in Kathmandu valley.

## Introduction

Avian influenza virus (AIV) type A strains are broadly classified into two categories based on pathogenicity: highly pathogenic avian influenza (HPAI), that causes severe illness and high mortality, and low pathogenic avian influenza (LPAI) that typically causes little or no clinical signs in birds [1]. Generally, HPAI is caused by AIV subtypes H5 or H7 but not all H5 and H7 are highly pathogenic [2]. HPAI has a zoonotic potential and can be transmitted to human from infected birds [1]. AI subtype H9 are generally but not always LPAI, as subtype H9N2 circulating in the Eurasian region has caused huge economic losses to the poultry industry, owing to decline in egg production and mortality when associated with other infections [3]. Also, as this virus has human like receptor specificity [4], it possess a potential to transmit to human posing a public health threat [5–6].

Nepal is an agrarian based economy and livestock sector including fisheries contributes nearly 12.5% to the total GDP. Among livestock sub-sector, poultry alone contributes nearly four percent of the GDP [7]. The total population of poultry birds in Nepal is estimated to be nearly 72 million [8]. During the last three decades, poultry industry globally has undergone rapid changes and shifting towards intensive production system, enhanced biosecurity, introductions of commercial breeds and application of preventive health measures [9]. While in developing countries like Nepal, these adoptions are limited due to high infrastructure cost for maintenance of biosecurity, quality hybrid chicks, qualitative feed, biologicals and quality veterinary care [10].

The booming poultry industry of Nepal has been hit by periodic outbreaks of avian influenza creating a great loss to poultry industry. Nepal recorded the first HPAI outbreak in eastern part of Nepal, Jhapa on January 16, 2009 where 28,000 poultry were killed to control the disease [11]. Thereafter, Nepal experienced several outbreaks in years 2010, 2011, 2012, 2013, 2017, 2018 and 2019 [12]. From August 2016 to July 2017, 3.85% (6/156) swab samples were positive for H5 and 30.13% (47/156) samples were positive to subtype H9 by Real Time Reverse Transcriptase Polymerase Chain reaction (RT-PCR) at CVL. During the same period, out of 3930 cloacal and tracheal swab samples collected for bio-surveillance, 0.41% (16/3930) samples were positive for H9. Likewise, from August 2017 to July 2018, 410 samples were received in CVL where 1.95% (8/410) samples were tested positive for H5N1 and 71.95% (295/410) were tested positive to H9. Out of 1597 swab samples collected for bio-surveillance, 6.9% (110/1597) were tested positive for H9. The molecular tests performed on samples submitted from Nepal at OIE reference lab, Australian Animal Health Lab (AAHAL), Australia identified H5N1 virus to be of clade 2.3.2.1a and H9N2 to be of G1-like H9N2 lineage with closest relationship to other G1-like H9N2 viruses that circulate in the South Asian region [13].

Kathmandu valley (Kathmandu, Bhaktapur and Lalitpur), the capital of Nepal has been identified as a high risk area for both LPAI and HPAI 14]. There have been several outbreaks of AI subtype H5 and H9 in Kathmandu valley since 2013 [13], which have caused massive economic loss and direct negative effect on the livelihood of the farmers. In addition, first human death case of AI subtype H5 was confirmed in Nepal in May 2019 [15]. Though there have been increase in the number of AI subtype H9 outbreaks, limited studies have been conducted to investigate the causes associated with these outbreaks. The identification of the potential risk factors would be helpful to mitigate the disease outbreaks in the future. The objective of this study is to identify the risk factors associated with AI subtype H9 outbreaks in Kathmandu valley.

## Materials and Methods

### Case definition and control farm selection

A retrospective case-control design was used in this study. The case registry book of Central Veterinary Laboratory (CVL), Tripureswor, Kathmandu was accessed from March 2018 to April 2019 for the study. A farm was considered as a case if it was confirmed positive for AI subtype H9 in rapid antigen detection test followed by Polymerase Chain Reaction (PCR). The control farm was any farm with no history of AI subtype H9 outbreak and are closer to the case farms (≤3 km from case farms). The ratio of case and control farm is 1:1 with 50 cases and 50 control farms. The distance between the case and the control farms was estimated using Google maps version 9.87.4.

Data were collected using a structured questionnaire having nineteen objective and open-ended questions. Questionnaire were pre-tested at ten farms of Kathmandu valley for its validity. There is no need for ethical review to collect questionnaire based data about the management of poultry farms in Nepal. Yet, the verbal consent was obtained from the farm owners. The preliminary interview was conducted to the poultry owners who came for the diagnostic services at CVL and subsequent farm visit was made to get detailed farm information. Some of the case farms and the control farms were contacted by phone to schedule the meeting for the interview to get more information about the outbreak and the biosecurity status.

The risk factors for the detection of Avian Influenza were identified from literature review and expert’s opinion [16]. The risk factors selected were broadly divided into following categories: i) Farm and bird characteristics ii) Farm management and ii) Biosecurity situation of the farm. In the farm and bird characteristic category, we documented farm location, farm type, age of farm, type of birds, age, flock size, number of flocks, and mortality patterns in the farm. In farm management category, the variable documented include interval between two batches of birds, fumigation of farms before introducing new batch, culling of birds during morbidity, flooring type, water source and previous history of AI outbreak. To assess biosecurity level, we documented use of aprons, boots, and self-sanitization before entering farm, presence of other animals and birds in farm, litter disposal, dead bird disposal, distance of nearby farm, type of nearby farm, vehicles allowed in farm, fencing and distance from main road.

### Site of study

Study was conducted in the poultry farms of Kathmandu valley. Kathmandu valley consists of three districts including capital city, Kathmandu and adjacent districts Lalitpur and Bhaktapur.

### Statistical analysis

Data were entered in Microsoft Excel 2016 and converted to CSV file for risk factor analysis in STATA 14.2. All continuous variables were transfigured into categorical variables using quartiles and averages to avoid problem of linearity. The 2×2 table and chi-square test was performed to test independence between variables using online software OpenEpi version 3.01 and corresponding p-values were calculated.

Univariable logistic regression analysis was applied to test association of individual risk factors with the detection of AI subtype H9. Odd ratios (ORs), their 95% confidence intervals (CIs) and corresponding p-values were estimated by logistic functions in STATA. Variables that met a cut-off of p≤ 0.2 in the univariable logistic regression were considered for the final multivariable logistic regression. The adjusted odds ratios from the multivariable regression were calculated to measure the strength of associations of the risk factors to detection of AI subtype H9 in poultry farms of Kathmandu valley. The fitness of the final multivariable model was evaluated using the “estat gof’ functions of Hosmer-Lemeshow test in STATA.

## Results

### Population characteristics

The epidemic curve of AI subtype H9 outbreaks on farms of Kathmandu valley from March 2018 to April 2019 is shown in Figure 1. There were altogether 105 farms detected positive to AI subtype H9 during the study period in Kathmandu Valley. An outbreak started from March 2018 and the highest number of cases were observed in May 2018 with 16 farms infected which gradually decreased to one case farm in September 2018. Again, in November 2018, the number of infected farms rose to 16 and the outbreaks continued until January 2019. Later in March 2019, the outbreaks boomed to 24 and on average eight farms remained infected until April 2019. Altogether 76 (61.9 %) commercial broilers, 30 (24.4%) layers, 14 (11.4%) backyard poultry (local chicken and duck) and three (2.44%) breeder farms were confirmed positive to H9 by PCR at the period of study. The mean flock size of the studied farms was 2018 (95% CI: 1686.16, 2350.04) and the median farm size was 1700 (Range: 12-15000).

**Fig. 1:**
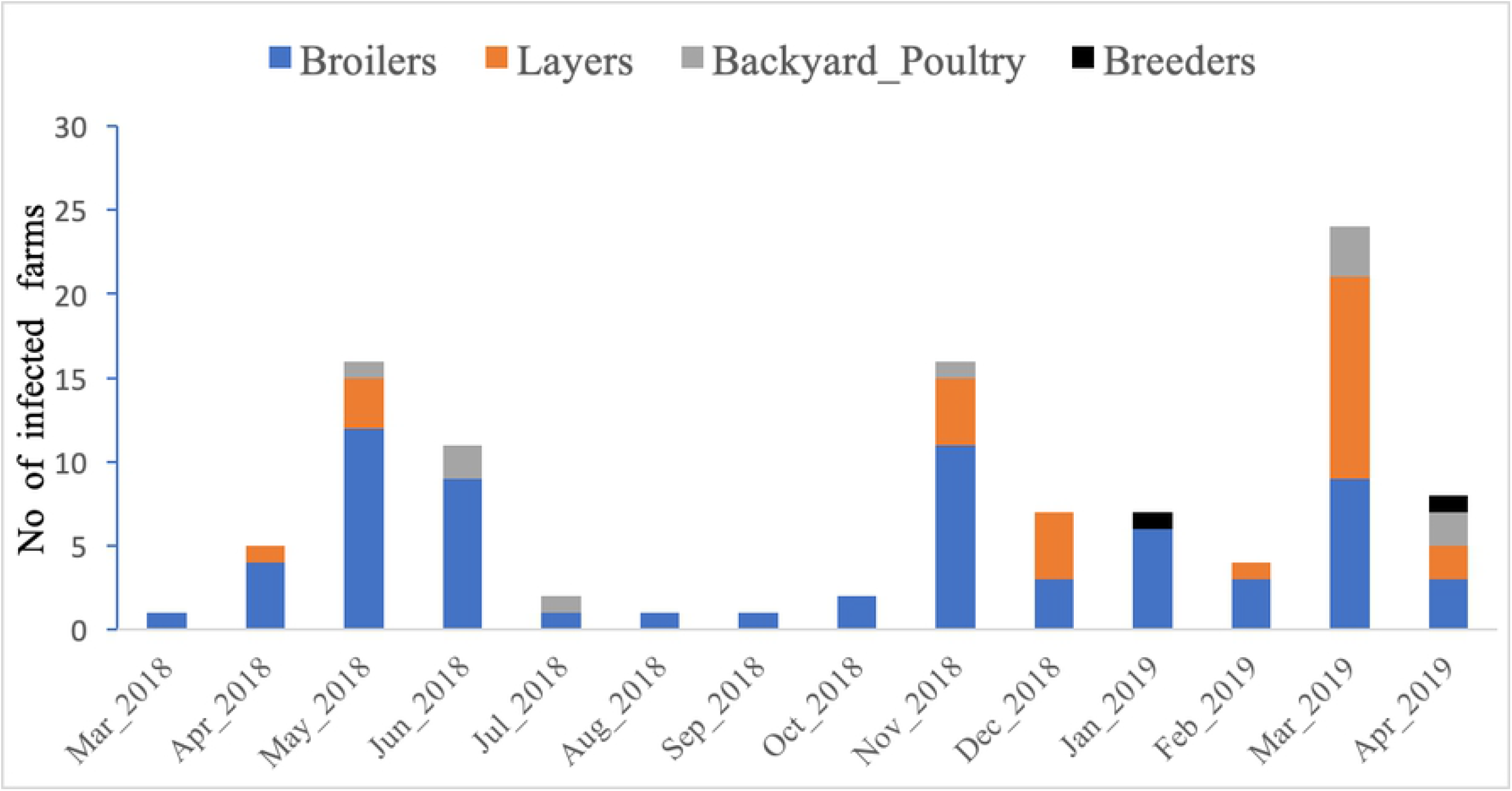
Epidemic curve for avian influenza subtype H9 infected farms in Kathmandu valley, Nepal

### Univariable analysis of risk factors

We selected total nineteen variables for the univariable analysis under three different categories. Under the bird and farm characteristics category: out of eight variables tested, six variables were significantly associated with the detection of AI subtype H9. Bird of ages between 31 to 40 days (OR= 4, 95% CI: 1.0-16.31), flock size of less than or equal to 2000 (OR= 2.9, 95% CI: 1.06-8.07), total mortality percentages of >30 to 50 (OR= 10.7, 95% CI: (1.22-93.92) and >50 to 80 (OR= 6.1, 95% CI: (1.17-31.92), the farm types of giriraj and kuroiler (backyard poultry) are significantly protective to H9 compared to commercial broiler (OR= 0.14, 95% CI: 0.02-1.18) (Table-1).

**Table 1:** Univariable logistic regression analysis of risk factors related to bird and farm characteristics

| Variables | Category | No of cases<br>(n=50) | No of controls<br>(n=50) | OR | 95% CI | P value |
| --- | --- | --- | --- | --- | --- | --- |
| Age of birds | 1-20 days | 5 | 10 |  |  |  |
|  | 21-30 days | 9 | 13 | 1.4 | (0.35-5.44) | 0.641 |
|  | 31-40 days | 14 | 7 | 4 | (1.0 -16.31) | 0.050* |
|  | 41 to 60 days | 6 | 4 | 3 | (0.57-15.77) | 0.194 |
|  | >60 | 16 | 16 | 2 | (0.56-7.18) | 0.288 |
| Flock size | Upto 2000 | 30 | 41 | 2.9 | (1.06-8.07) | 0.038** |
|  | >2000 to 5000 | 15 | 7 | 3.4 | (0.62-18.82) | 0.158 |
|  | >5000 | 5 | 2 |  |  |  |
| No of Poultry Shed | >3 | 5 | 1 | 5.4 | (0.61-48.40) | 0.128 |
|  | <=3 | 45 | 49 |  |  |  |
| Total mortality<br>percentage (%) | <=10 | 19 | 29 |  |  |  |
|  | >10 to 30 | 12 | 18 | 1.0 | (0.40-2.58) | 0.971 |
|  | >30 to 50 | 7 | 1 | 10.7 | (1.22-93.92) | 0.033** |
|  | >50 to 80 | 8 | 2 | 6.1 | (1.17-31.92) | 0.032** |
|  | >80 | 4 | 0 | 1.0 |  |  |
| Age of farm | <=3 years | 34 | 41 |  |  |  |
|  | >3 to 5 years | 7 | 5 | 1.69 | (0.49-5.80) | 0.406 |
|  | >5 years | 9 | 4 | 2.71 | (0.77-9.59) | 0.121 |
| Type of farm | Commercial Broiler | 27 | 30 |  |  |  |
|  | Commercial Layers | 15 | 9 | 1.85 | (0.70-4.92) | 0.216 |
|  | Breeder | 1 | 0 | 1 |  |  |
|  | Duck | 2 | 0 | 1 |  |  |
|  | Giriraj_and_kuroiler | 1 | 8 | 0.14 | (0.02-1.18) | 0.071 |
|  | Local_backyard | 4 | 3 | 1.48 | (0.30-7.23) | 0.627 |
| Nearby poultry farm<br>located within a<br>distance | <500 meter | 35 | 31 | 1.43 | (0.62-3.29) | 0.399 |
|  | >=500 meter | 15 | 19 |  |  |  |
| Farm distance from the<br>main road | <=1km | 45 | 44 | 1.23 | (0.35-4.32) | 0.75 |
|  | >1km | 5 | 6 |  |  |  |

\*\* P – value <0.05, statistically significant
CI- Confidence interval; OR- Odds ratio

Among the five variables under the farm management category, three variables were significantly associated with the AI subtype H9 outbreak. The associated variables were: “no fumigation “(OR= 2.8, 95 % CI: 1.11-7.01), “no culling of sick birds” (OR= 1.10, 95 %CI: 0.46-2.62), “water supply by boring compared to tanker supply” (OR= 3.4, 95 % CI: 0.90-13.26) and “the previous history of AI outbreak” (OR= 7.98, 95 %CI: (0.94-67.46) (Table 2).

**Table 2:**
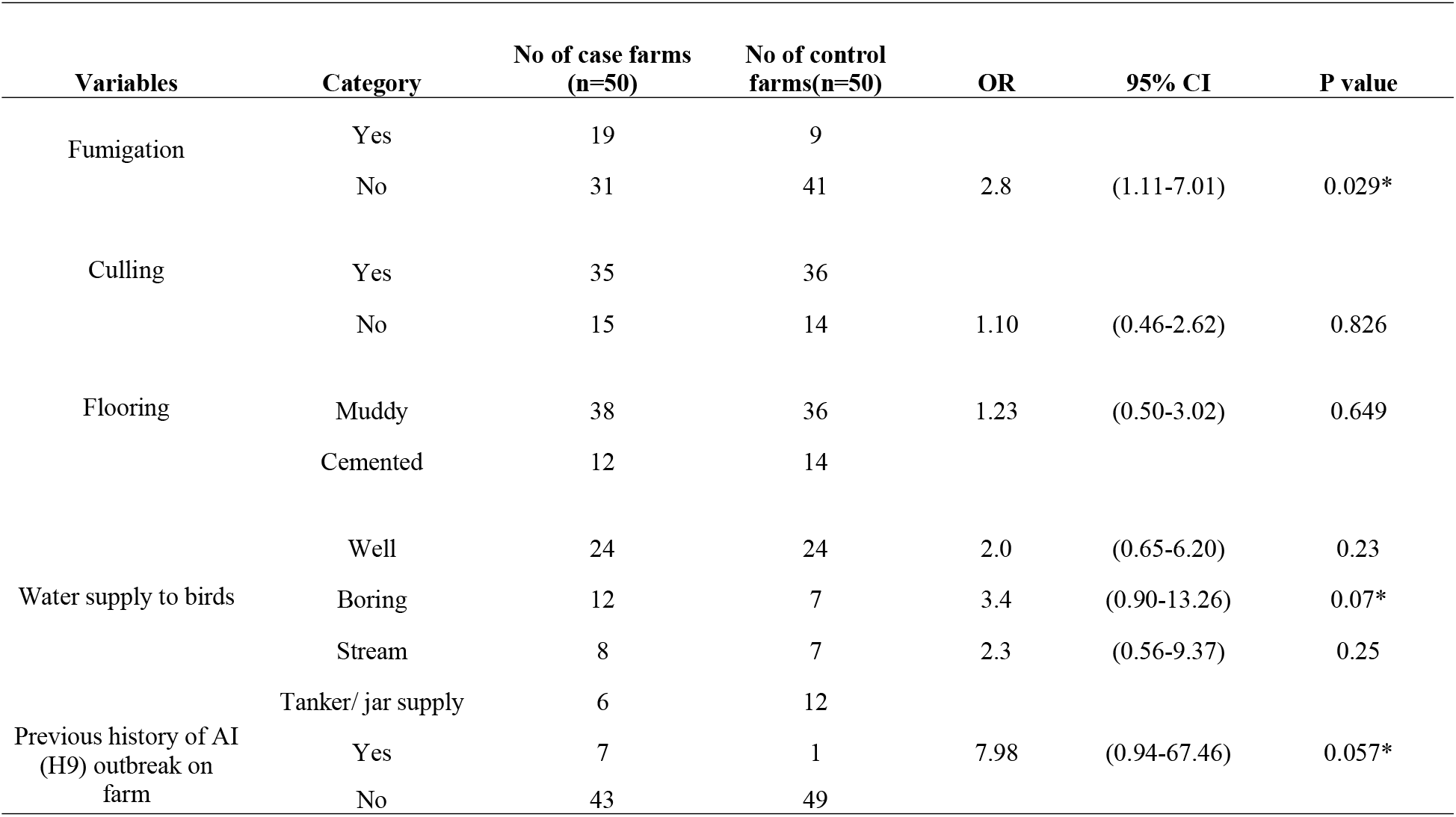
Univariable logistic regression analysis of risk factors related to farm management

In the biosecurity category, six risk factors were identified and two risk factors that were significantly associated with were: “no boots application while entering farm “(OR= 2.4, 95 % CI: 0.98-5.68) and “no foot bath at entry of farm “(OR= 3.32, 95 % CI: 1.36-8.09) (Table 3).

**Table 3:**
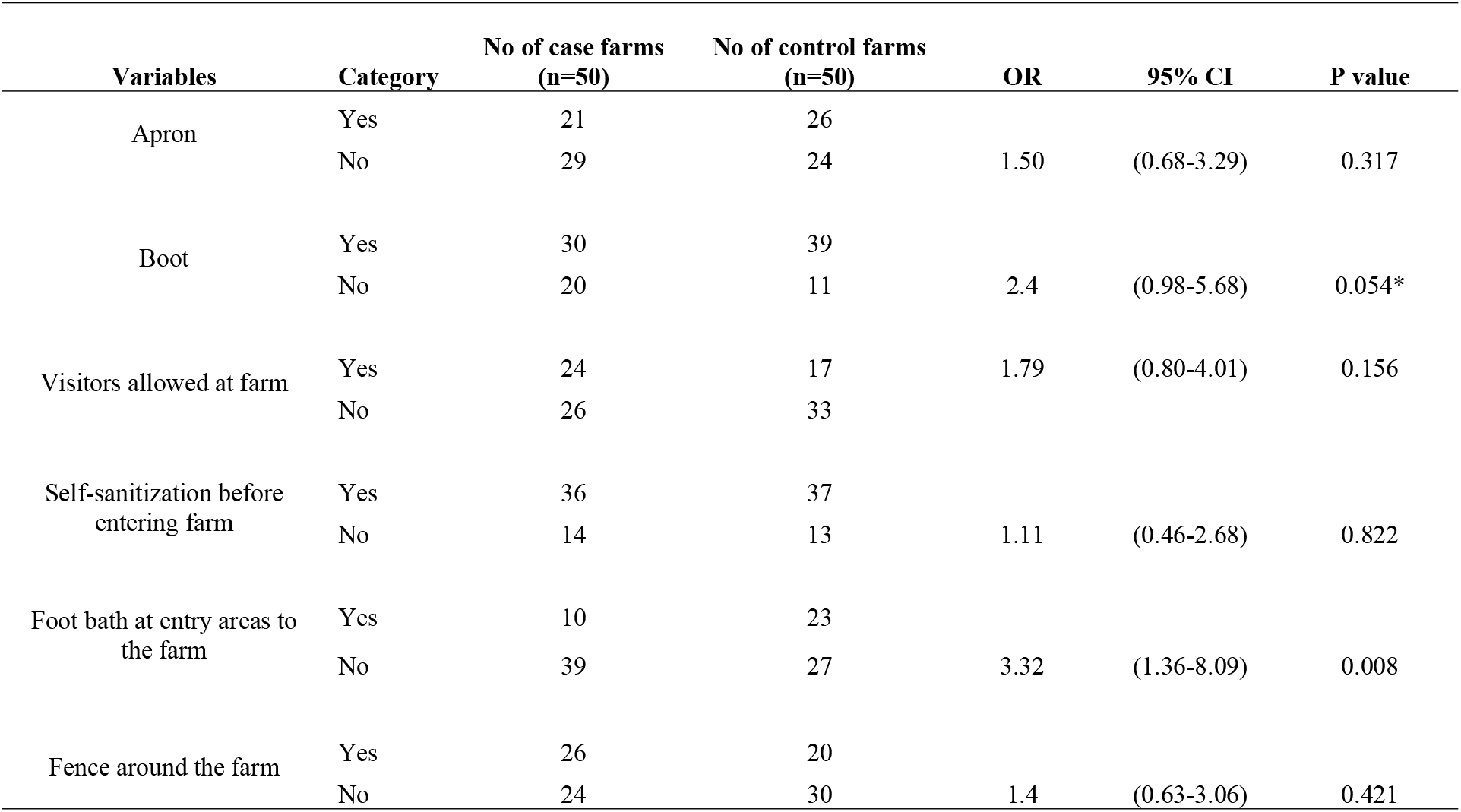
Univariable logistic regression analysis of risk factors related to farm biosecurity

### Multivariable logistic regression analysis

Altogether fifteen variables: six from bird and farm characteristics (Table-1), five from the farm management status category (Table-2) and three from the farm biosecurity status category (Table-3) were included in the multivariable logistic regression based on the cut-off criteria.

Ten factors were identified as the risk factors in the final model Table 4. The birds of age category from 31 to 40 days were 11 times more likely to be tested positive to AI subtype H9 compared to birds of age category 1 to 20 days (OR= 11.31, 95% CI: 1.31-98.02) (p=0.028) keeping others variables constant. The total mortality percentage of birds’ due to the AI subtype H9 were more likely between the range of 30 to 50 percentage (OR= 144.7, 95% CI: 4.53-4622.49) (p=0.005).

**Table 4:**
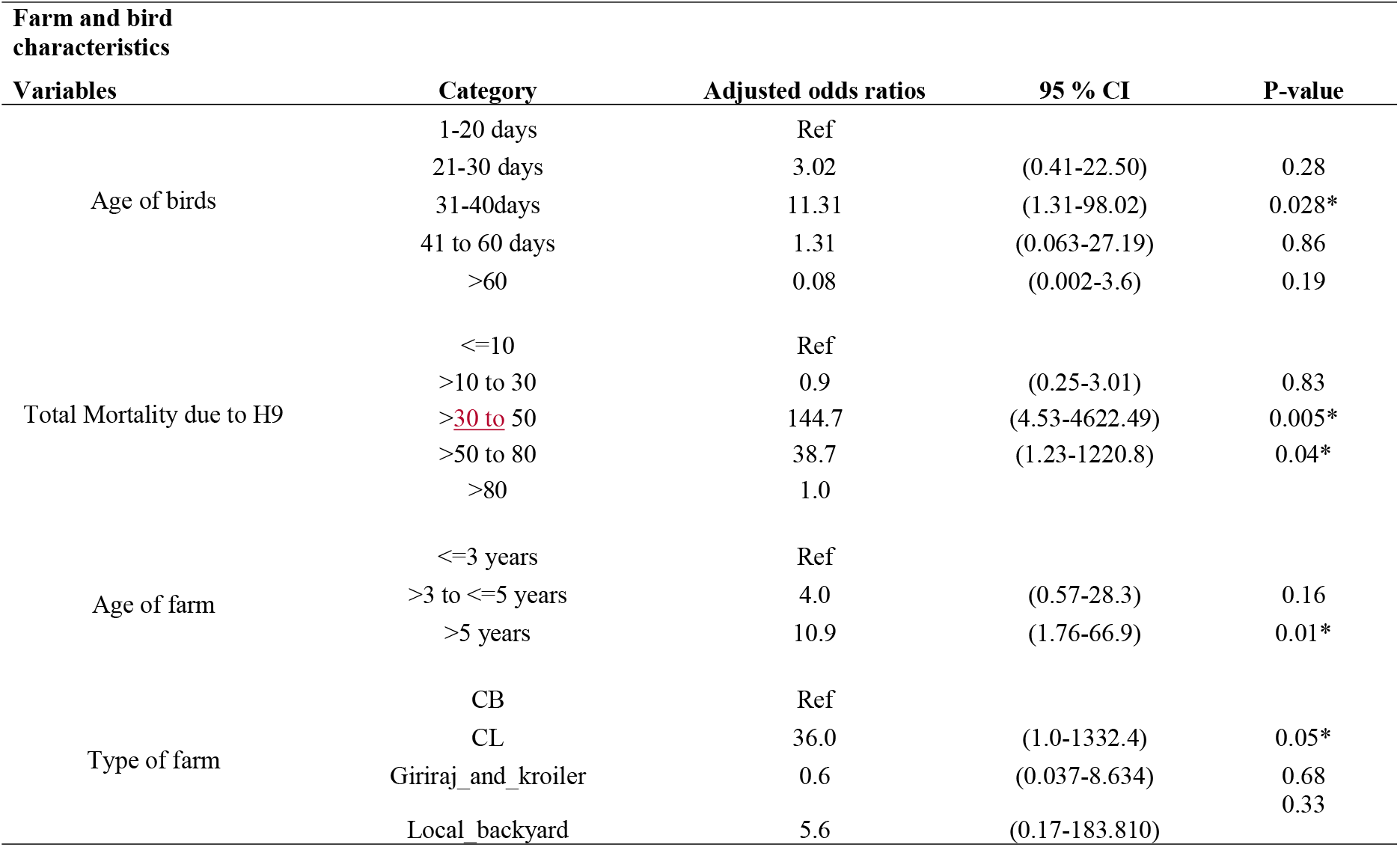
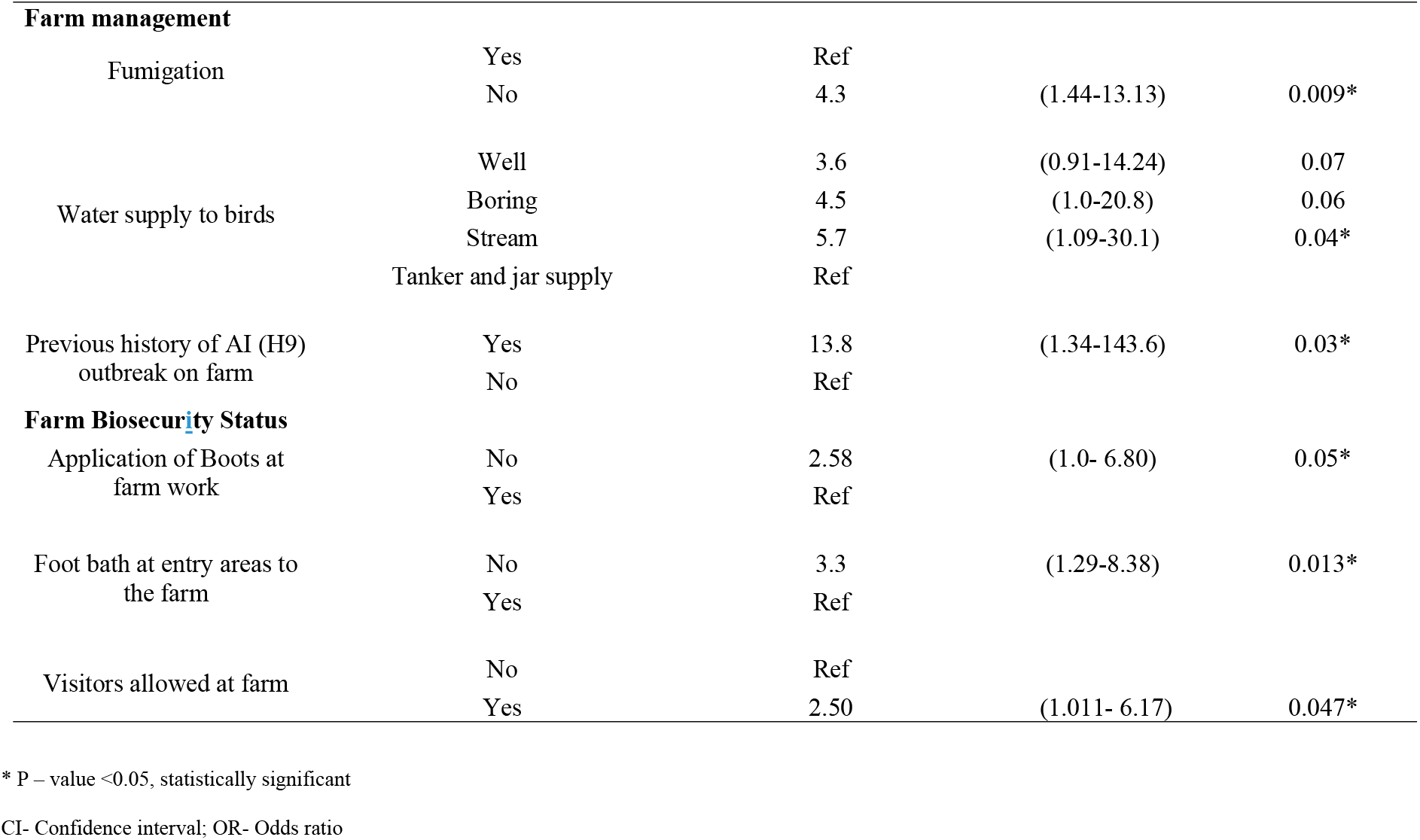
Multivariable logistic regression analysis related to farm and bird characteristics, farm management and biosecurity status

The farms older than five years were almost eleven times more likely to be detected for avian influenza (OR= 10.9, 95% CI: 1.76-66.93) compared to farms up to three years (p=0.01). The commercial layers farms are thirty-six times (OR= 36.0, 95% CI: 1.0-1332.40) more likely to be detected with the AI subtype H9 compared to commercial broilers farms (p=0.052).

The farms house that were not fumigated for every batch of poultry were four times (OR= 4,95 %CI: 1.44-13.13) more likely to be tested positive to AI subtype H9 compared to farms that got fumigated (p=0.009). In the water supply process on farms, the farms that used stream water for feeding poultry birds were almost significantly six times (OR= 5.7, 95% CI: 1.10-30.13) in risk of detecting AI subtype H9 compared to farms that supplied city water tank respectively (Table 4). The farms that had a previous history of AI outbreak were almost fourteen times (OR= 13.8, 95 %CI: 1.34-143.63) more likely to be detected with AI subtype H9 (p = 0.028).

In the biosecurity status “that farms that did not applied boots to visitors while entering the farms were 2.58 times “(OR= 2.58, 95% CI: 0.98-6.80) at risk of detecting AI subtype H9 compared to farms that applied boots but the result is borderline significant (p=0.055). The farms that allow visitors to enter farm are almost 2. 5 times (OR= 2.5, 95% CI: 1.011, 6.17) more likely in detecting the AI subtype H9 compared to farms that did not allow visitors to enter the farm (p=0.047). The farms that had no foot bath at entrance are almost three times (OR= 3.3, 95 %CI: 1.29, 8.38) more at risk of detecting AI subtype H9 compared to farms that had no foot bath at entrance (p=0.013).

## Discussion

This is the first case-control study conducted in Nepal to identify the risk factors associated with AI subtype H9 outbreaks in Nepal to the best of our knowledge. The results indicated that several farm and bird characteristics, farm management and biosecurity situations of the farms were associated to the detection of AI subtype H9 in poultry farms of Kathmandu valley.

Under the bird’s characteristics category, the birds of ages between 31 to 40 days are highest risk of contracting AI subtype H9 indicating special attention should be given to the birds during that age by the farmers. A study in Pakistan [17] found that the “age of flock at the time of submission of samples >50 days” was a risk factor associated with outbreak of AI subtype H9N2 in commercial poultry farms of Pakistan. The mortality percentage due to disease was as high as 80 percentage in case farms in the birds. It may be probably due to secondary infections such as new castle disease and infectious bursal disease and E coli [17]. The farm-house that was greater than five years old were at higher risk of detecting AI subtype H9. As the farm production system becomes older, the biosecurity facilities may become older and disrupted. Also, as the farms grows older, it keeps producing many batches of poultry such that there is higher burden of virus around the poultry surroundings [16]. Also, the commercial layers were at higher risk of detecting AI subtype H9. The probable reason could be the poor biosecurity status of the poultry farms such as movement and exchange of old egg trays between which is commonly practiced in Nepal.

Under the farm management category, the farms house that were not fumigated for every batch of poultry were four times more likely to be tested positive to AI subtype H9 compared to farms that got fumigated. This may be due to bacteria and virus that are missed by regular disinfection can be destroyed by fumigation only. The farms that use stream water for feeding poultry birds are significantly six times at risks of detecting AI subtype H9 compared to farms that used bulk tank water supply. The stream water is source of environmental water where the wild birds that acts as mechanical source for contaminating water by their droppings [9,18–19]. The odds of AI subtype H9 outbreak is almost fourteen-fold greater for a farm that have previous history of AI outbreak than those without history of AI outbreak. In the biosecurity status, “that farms that did not applied boots to visitors while entering the farms were 2.58 times at risk of detecting AI subtype H9 compared to farms that applied boots. This finding is nearly consistent to study by Chaudhary et al., 2015, where “worker change disinfected boots” was found as risk factor associated with outbreak of AI subtype H9N2 in commercial poultry farms of Pakistan. The farms that allow visitors are almost 2.5 times more likely in detecting the AI subtype H9 compared to farms that did not allow. The farms that had no foot bath at entrance are almost three times more at risk of detecting AI subtype H9 compared to farms that had no foot bath at entrance. This result is consistent with the study conducted in south Korea [16].

In this study, some of the variables such as flock size, number of poultry sheds in the farm, and use of aprons during the farm operations are not significantly associated with detection of AI subtype H9 which was found to be significantly associated with detection of AI subtype H9 in other studies, which may be either due to difference in the poultry productions system of Nepal from other countries or limited number of observations for the case and control and control data.

## Limitations of the study

The number of cases and the control farms selected is lower as many of the farms and owners were not reachable at the time of study such that the level of significance for some variables are not achieved or are with the wider confidence interval of ORs. The farms that are close to CVL are more likely to submit samples than the farms located far away from the laboratory leading to selection bias. The farmers who are aware of AI and the diagnostic capability of the laboratory are more likely to visit the laboratory for the confirmation of the disease and small farms might have been missed. Some farms could not answer some questions such as “do you fumigate your farm?” as some of them are not aware of term fumigation that may lead to response bias. Also, some breeder farms were not willing to disclose their previous history of AI as they were paranoid of rejecting the chicks of their hatchery by the dealer if they know that their parent birds were AI infected.

## Conclusion

We identified risk factors related to poultry bird characteristics, farm management and farm biosecurity characteristics that contributes to outbreak of avian influenza AI subtype H9 among the poultry farms of Kathmandu Valley. The study pinpoints importance of good management and application of strict biosecurity measures for the control of AI subtype H9 outbreaks in the poultry farms. This study can be a baseline for similar studies in future.

## Recommendation

Good management and strict biosecurity can prevent AI subtype H9 infection in Kathmandu valley. Management of identified risk factors is a key consideration to mitigate the future risks of AI subtype H9 outbreak in Kathmandu valley. We suggest more detailed analytic study in the future.

## Supporting Information

**S1 Appendix.** Questionnaire for “Risk factors associated with AI subtype H9 outbreaks on poultry farms in Kathmandu valley, Nepal” (PDF).

## Acknowledgement

I want to acknowledge all the poultry farmers of Kathmandu valley who warmly welcomed us on their farms and participated actively in answering the questions to our research.

## Authors contributions

Conceived and designed the experiments: TRG. Performed experiments: TRG, BRS, Analyzed the data: TRG. Performed laboratory procedures and contributed reagents/materials/analysis tool: DDB, PK, MM and TRG. Wrote first draft of the paper: TRG. Edited the first draft and contributed to finalize the paper: SK and TRG.

